# CNVeil enables accurate and robust tumor subclone identification and copy number estimation from single-cell DNA sequencing data

**DOI:** 10.1101/2024.02.21.581409

**Authors:** Weiman Yuan, Can Luo, Yunfei Hu, Liting Zhang, Zihang Wen, Yichen Henrry Liu, Xian Mallory, Xin Maizie Zhou

**Author notes:** Equal contributor (Weiman Yuan, Can Luo).

## Abstract

Single-cell DNA sequencing (scDNA-seq) has significantly advanced cancer research by enabling precise detection of chromosomal aberrations, such as copy number variations (CNVs), at a single-cell level. These variations are crucial for understanding tumor progression and heterogeneity among tumor subclones. However, accurate CNV inference in scDNA-seq has been constrained by several factors, including low coverage, sequencing errors, and data variability. To address these challenges, we introduce CNVeil, a robust quantitative algorithm designed to accurately reveal CNV profiles while overcoming the inherent noise and bias in scDNA-seq data. CNVeil incorporates a unique bias correction method using normal cell profiles identified by a PCA-based Gini coefficient, effectively mitigating sequencing bias. Subsequently, a multi-level hierarchical clustering, based on selected highly variable bins, is employed to initially identify coarse subclones for robust ploidy estimation and further identify fine subclones for segmentation. To infer the CNV segmentation landscape, a novel change rate-based across-cell breakpoint identification approach is specifically designed to diminish the effects of low coverage and data variability on a per-cell basis. Finally, a consensus segmentation is utilized to further standardize read depth for the inference of the final CNV profile. In comprehensive benchmarking experiments, where we compared CNVeil with seven state-of-the-art CNV detection tools, CNVeil exhibited exceptional performance across a diverse set of simulated and real scDNA-seq data in cancer genomics. CNVeil excelled in subclone identification, segmentation, and CNV profiling. In light of these results, we anticipate that CNVeil will significantly contribute to single-cell CNV analysis, offering enhanced insights into chromosomal aberrations and genomic complexity.

## Introduction

Copy number variations (CNVs) are variations that alter the number of copies of genomic regions. They are frequent somatic mutations that play a crucial role in cancer as well as a variety of other genetic diseases [1–4]. These variations manifest themselves not only within primary tumor sites but also during metastasis [3,5]. This phenomenon poses challenges to standardized treatment and emphasizes the need for tailored therapeutic strategies [6]. Customizing treatments based on subclone structure shows potential in mitigating the risk of cancer recurrence, as it addresses not only the predominant pathogenic gene alterations [7–10]. Further advancement of tumor treatment necessitates a more sophisticated cellular analysis to discern and comprehend the nuances of tumor diversity.

Single-cell DNA sequencing (scDNA-seq) provides a detailed view of the genome at the individual cell level [11, 12], enhancing the comprehension of a tumor’s clonal architecture. This biotechnological approach, by individually analyzing targeted cells, facilitates a more precise analysis of CNVs and captures the pathogenic evolution of tumor subclones [13, 14]. In contrast to bulk sequencing, which characterizes the genomic landscape at the population level, scDNA-seq avoids the averaging effect that could obscure distinctive CNV profiles [15–17]. While single-cell RNA sequencing (scRNA-seq) primarily offers insights into the expressed regions of the genome rather than its entirety, scDNA-seq investigates the complete genome, offering an accurate depiction of CNVs [18–21]. However, scDNA-seq is vulnerable to sequencing errors and biases that may introduce distortions to the data. These errors, coupled with the typically low and nonuniform depth of coverage per cell, which is induced by the non-linear amplification and dropout events during the library preparation and sequencing procedures [4, 22], can impede the detection of CNVs. This underscores the imperative for meticulous approaches to accurately unveil CNV profiles.

To date, several methods have been developed to investigate single-cell CNVs from scDNA-seq data. HMMcopy [23] was initially introduced to detect CNVs from the bulk sample and is also applicable for scDNA-seq data. It employs an eleven-state Hidden Markov Model (HMM) that incorporates GC content correction to mitigate false positives. Similarly, Ginkgo [24] addresses data bias by examining the relationship between GC content and read depth. Developed in 2015, it stands out as the first CNV inference tool explicitly designed for scDNA-seq data, utilizing circular binary segmentation (CBS) for CNV detection. The following year, AneuFinder [25] also leveraged an HMM for CNV detection in scDNA-seq data, coupled with quality control through multivariate clustering. These methods, either employing the HMM or CBS algorithm, focus on segmenting the genome at the individual cell level. Between 2020 and 2022, several multiple-cell-based methods, including SCOPE [19], CHISEL [26], and SeCNV [27], have become available. These methods exploit the principle that cells from the same subclone are likely to share common CNV breakpoints, enabling the inference of CNVs by overcoming the challenges posed by low and non-uniform coverage at the single-cell level. Multiple-cell-based methods handle the scDNA-seq data more effectively by leveraging shared information among individual cells. Theoretically, multiple-cell-based methods are expected to achieve better performance than single-cell-based methods. SCOPE clusters cells through a generalized likelihood ratio test and optimizes the segmentation count using a modified Bayesian Information Criterion (BIC). SeCNV partitions the genome into segments by minimizing the structural entropy from a depth congruent map. Both SCOPE and SeCNV incorporate strategies for identifying normal cells within noisy datasets to establish a baseline for bias correction; SCOPE utilizes the Gini coefficient, while SeCNV utilizes the coefficient of variation (CV). CHISEL advances allele-specific CNV detection, incorporating phasing and Expectation-Maximization algorithms to address high allelic dropout issues. Recently, a deep learning-based convolutional autoencoder framework, rcCAE [28], was introduced for noise reduction and genome segmentation, aiming to infer CNV in scDNA-seq data.

Although most existing methods for inferring CNV from scDNA-seq data show promise in certain scenarios, their performance is not consistently robust across all cells. Often, these methods are sensitive to data variability induced by the complexity inherent in cancer data. To address these challenges, we introduce CNVeil, a robust quantitative algorithm designed to accurately reveal CNV profiles while overcoming the inherent noise and bias in scDNA-seq data. CNVeil offers several innovative and beneficial features and performance. (1) It implements a PCA-based Gini coefficient to select normal cells as normal controls to normalize the read depth. (2) CNVeil defines and selects highly variable bins as a feature vector to classify normal and tumor cells. (3) It performs different levels of clustering to either estimate optimal ploidy or conduct cross-cell breakpoint and segmentation identification. (4) CNVeil standardizes the read depth by utilizing cross-cell segmentation to maintain genuine genomic aberrations while minimizing the impact of artifact noise. (5) It achieves better subclone identification, segmentation, and CNV profiling across all datasets. (6) It shows robust performance across all cells and bins.

## Results

### CNVeil revealed CNV profile reliably on simulated datasets

To evaluate CNVeil and benchmark existing tools, the lack of ground truth data as a gold standard for most datasets is the most challenging issue. We thus first employed SimSCSnTree [29] to simulate datasets, providing a gold standard. SimSCSnTree stochastically simulates a tree and imputes CNAs on the tree. Specifically, SimSCSnTree allows users to decide multiple factors such as ploidy of the cells, tree structure, number of clones, number of cells, size of the CNAs, ratio between amplification and deletion, whether there is whole genome duplication, and so on. For the experimental design of our simulations, we generated four different datasets by tuning the average ploidy varying from 1.5 to 5. The number of tumor cells in each dataset was 96, 97, 95, and 100 respectively (Table 1). In addition, we also simulated 50 normal cells as negative control, intermixed with tumor cells as input for all CNV profiling tools. More simulation details are outlined in the methods section.

**Table 1:** Resource for different tools, datasets, and the gold standard. Top panel: The CNV inference methods used in this paper. The tool version number, cited article, and tool links are shown in the table. Bottom panels: The real and simulated scDNA-seq datasets and the aCGH gold standard used in this paper. The number of cells, identifiers, and links for each dataset are shown in the table. For simulated datasets, the corresponding simulator is listed.

| CNV callers | Version | Resource | Link |
| --- | --- | --- | --- |
| CNVeil | default | This paper | https://github.com/maiziezhoulab/CNVeil |
| rcCAE | default | Yu et al. 2023 | https://github.com/zhyu-lab/rccae |
| SeCNV | default | Wang et al. 2022 | https://github.com/deepomicslab/SeCNV |
| CHISEL | 1.0.0 | Zaccaria et al. 2019 | https://github.com/raphael-group/chisel |
| SCOPE | 1.10.0 | Wang et al. 2019 | https://github.com/rujinwang/SCOPE |
| AneuFinder | 1.27.1 | Bakker et al. 2016 | https://github.com/ataudt/aneufinder |
| Ginkgo | default | Garvin 2015 | https://github.com/robertaboukhalil/ginkgo |
| HMMcopy | 1.40.0 | Shah et al. 2006 | https://github.com/shahcompbio/HMMcopy |

| Dataset | Number of Cells | Identifier | Link |
| --- | --- | --- | --- |
| T10 | 100 | SRA: SRA018951 | https://www.ncbi.nlm.nih.gov/sra/SRX021401[accn] |
| T16P | 52 | SRA: SRA018951 | https://www.ncbi.nlm.nih.gov/sra/SRX037035[accn] |
| T16M | 48 | SRA: SRA018951 | https://www.ncbi.nlm.nih.gov/sra/SRX037132[accn] |
| KTN302 (pre-treatment) | 47 | SRA: SRP114962 | https://www.ncbi.nlm.nih.gov/sra/SRX3069874[accn] |
| KTN302 (mid-treatment) | 45 | SRA: SRP114962 | https://www.ncbi.nlm.nih.gov/sra/SRX3069873[accn] |
| Simulation<br>(ploidy = 1.5, 2, 3, 4, 5) | [96, 50, 97 95, 100] |  | https://github.com/compbiofan/SimSCSnTree |

Table 1: **Resource for different tools, datasets, and the gold standard.**
| Gold Standard | Number of Sectors | Identifier | Link |
| --- | --- | --- | --- |
| aCGH of breast cancer | 14 | GEO: GSE16607 | https://www.ncbi.nlm.nih.gov/geo/query/acc.cgi?acc=GSE16607 |

When assessing the CNV profile generated by each tool against the gold standard, we employed two different evaluation modes: segmentation mode and CNV mode. In segmentation mode, our comparison focused solely on the position of breakpoints (segmentation boundaries). If a tool accurately identified the break-point where the copy number changed, it was considered a correct breakpoint. In stringent CNV mode, an inference was regarded as correct if the tool identified both the correct breakpoint and the correct copy number for the left and right bins of the breakpoint. The evaluation was specifically applied to tumor cells. More details on the evaluation are also outlined in the methods section.

Overall, CNVeil demonstrated superior and robust performance when compared to the other six tools, rcCAE, SeCNV, SCOPE, AneuFinder, Ginkgo, and HMMcopy, across all four datasets. CHISEL was excluded from this analysis since SNP information was not incorporated into the simulated datasets. Specifically in the violin plot (Figure 2), CNVeil consistently achieved the highest average recall, precision, and F1 across various conditions, while maintaining a relatively low variation across all cells. In an exceptional condition where the dataset’s ploidy was 1.5, SCOPE had the best recall for segmentation, followed by CNVeil. Notably, we observed that existing tools exhibited significantly lower recall or higher variance in both segmentation and CNV evaluation when dealing with datasets featuring extremely low (1.5) and high (5.0) ploidy. In contrast, CNVeil maintained a high and robust performance, demonstrating its ability to adapt to changes in dataset ploidy. This superior ability was due to the fact that CNVeil employed a novel PCA-based Gini coefficient for identifying normal cells to correct bias and conducted an initial clustering of normal and tumor cells based on highly variable bins. CNVeil precisely detected all 50 normal cells through the PCA-based Gini coefficient procedure and the initial clustering step (Table S1).

**Figure 1:**
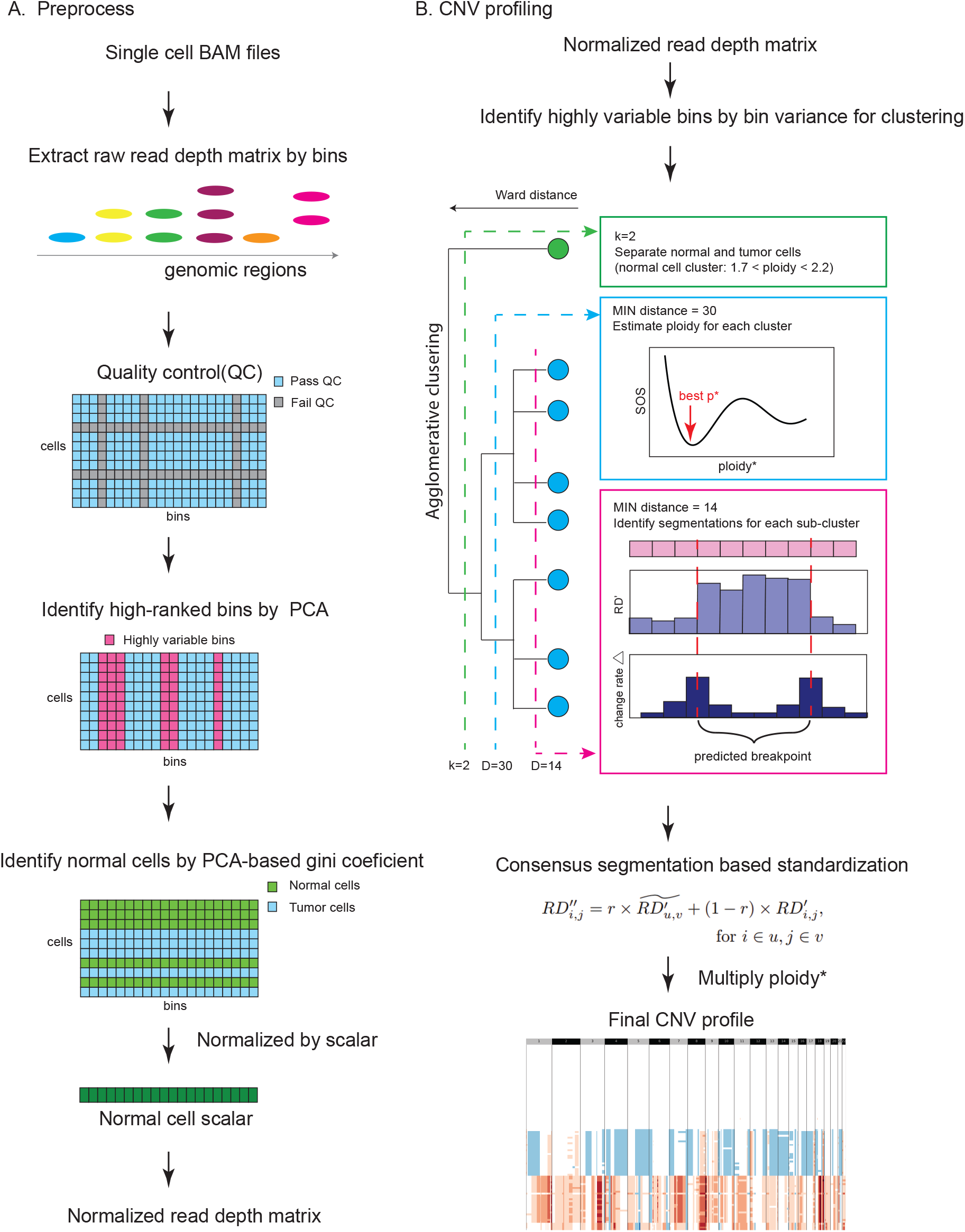
CNVeil workflow. The main workflow of CNVeil consists of six interconnected and conceptual modules: 1) Read count matrix construction (Preprocessing); 2) Data normalization by noise reduction and bias correction (Preprocessing); 3) Initial normal-tumor cell classification (CNV profiling); 4) Tumor sub-clones identification and ploidy estimation (CNV profiling); 5) Fine clustering and across-cell breakpoint and segmentation identification (CNV profiling); 6 Infer final CNV states (CNV profiling). A detailed description for each module is available in the method section.

**Figure 2:**
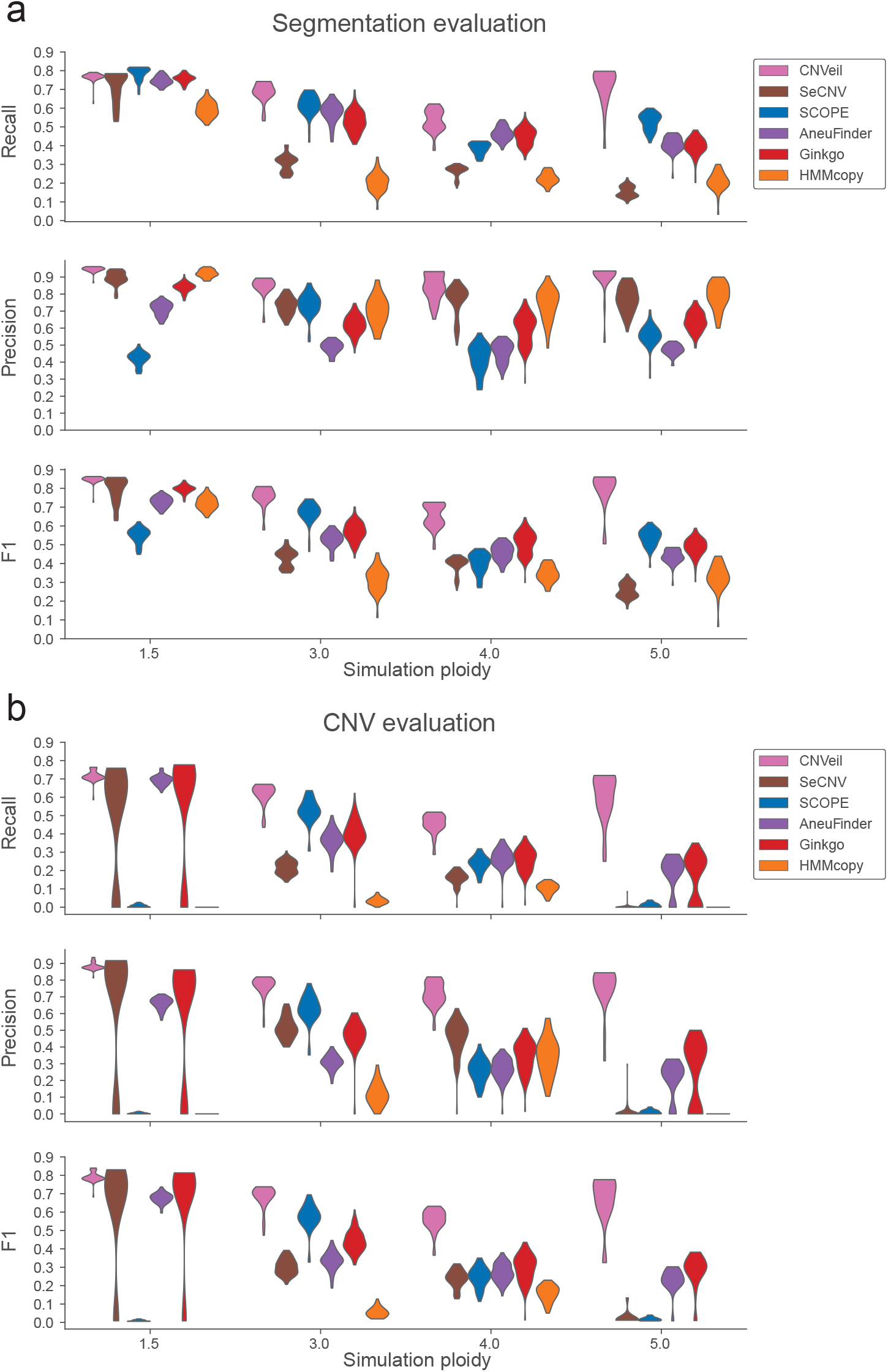
Segmentation and CNV evaluation against the gold standard in simulated datasets. **(a)** Violin plots illustrating recall, precision, and F1 scores for segmentation evaluation using six tools across four simulated datasets with varying ploidy values: 1.5, 3.0, 4.0, and 5.0. **(b)** Violin plots illustrating recall, precision, and F1 scores for CNV evaluation using six tools across four simulated datasets with varying ploidy values: 1.5, 3.0, 4.0, and 5.0. Tools are shown in chronological order by publication year.

We further used heatmaps to demonstrate the estimated copy numbers across all cells in all seven tools for each dataset. To ensure a fair and transparent comparison, we systematically ordered cells based on the gold standard. In the extremely high ploidy dataset (Figure 3), CNVeil maintained a highly similar CNV profile with the gold standard, while the other six tools either underestimated the ploidy of a significant number of tumor cells or did not infer correct copy numbers in many bins across the genome. In the extremely low ploidy dataset (Figure S1), CNVeil still maintained a highly similar CNV profile with the gold standard, however, other tools tended to overestimate the ploidy of a significant number of tumor or normal cells. In terms of the other two datasets (ploidy = 3 and 4.0, Figure S2-S3), CNVeil consistently generated the most comparable CNV profile to the gold standard, followed by SCOPE, Ginkgo, SeCNV, and AneuFinder.

**Figure 3:**
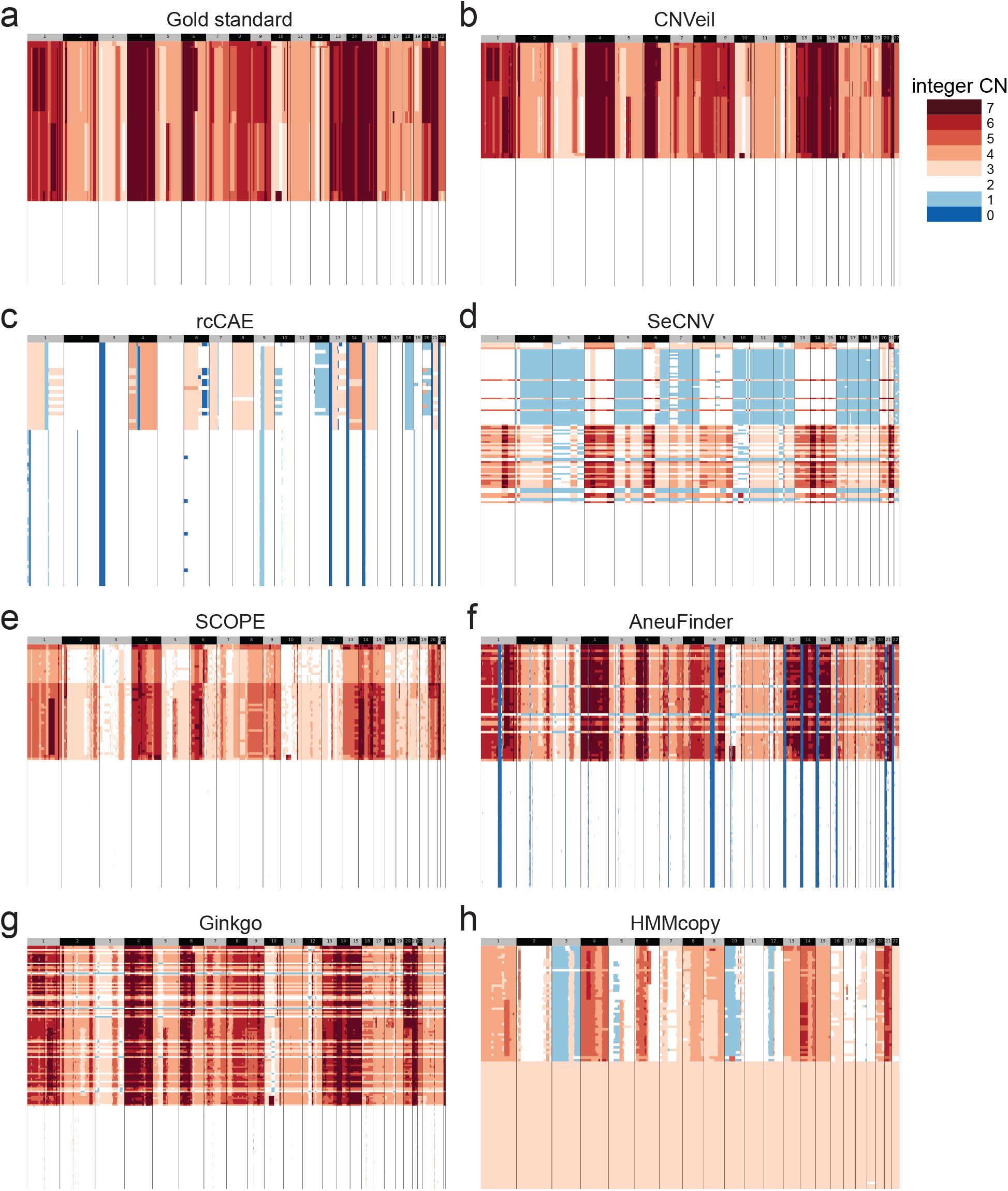
Comparison of inferred copy number profiles of single cells from the simulated dataset with a ploidy of 5.0. **(a)** Established copy number profiles by the gold standard. **(b-h)** Inferred copy number profiles by CNVeil, rcCAE, SeCNV, SCOPE, AneuFinder, Ginkgo, and HMMcopy. We ordered all heatmaps in a consistent cell order aligned with the gold standard which includes a normal cell subclone and a tumor subclone. Tools are shown in chronological order by publication year.

### Evaluation on the scDNA-seq data of breast cancer patient T10

We next benchmarked CNVeil and seven existing tools using real scDNA-seq data obtained from a breast cancer patient identified as T10 [30]. The dataset for patient T10 comprised 100 single cells in total (Table 1). Specifically, this dataset incorporated corresponding fluorescence-activated cell sorting (FACS) of the single cell data [31], which revealed four distinct cell subclones: A1 (hyperdiploid, ploidy = 2.85), A2 (hyperdiploid, ploidy = 3.1), H (hypodiploid, ploidy = 1.7), and D (diploid, ploidy = 2). FACS thus suggested a polygenomic tumor in T10.

The heatmaps in Figure 4 illustrate the estimated copy numbers across all cells in all eight tools. To ensure a fair and transparent comparison, we ordered all heatmaps in a consistent cell order based on the order of reported subclones in FACS. We observed that CNVeil, SeCNV, SCOPE, AneuFinder, and Ginkgo exhibited a generally similar CNV profile pattern. These five tools all identified a normal cell subclone, one subclone of hypodiploid cancer cells, and two subclones of hyperdiploid cancer cells, in order, in accordance with the FACS data. Specifically for the normal cell identification, CNVeil successfully identified 41 normal cells by the PCA-based Gini coefficient and 45 out of 45 gold normal cells through the initial clustering (Table S1). Upon closer examination, SeCNV, SCOPE, AneuFinder, and Ginkgo identified a few diploid cells as either hypodiploid or hyperdiploid. Additionally, SeCNV, AneuFinder, and Ginkgo identified a few hyperdiploid cells with unusually high ploidy levels. AneuFinder and Ginkgo also exhibited a few outlier bins that were not appropriately processed either during the preprocessing step or in subsequent stages. Among these five tools, CNVeil presented a clearer and more accurate CNV profile. We also observed that the copy number profiles of the two hyperdiploid subpopulations were remarkably similar, suggesting that the relapse originated from the same subclone present in the primary tumor. HMMcopy also identified four subclones, however, it tended to overestimate the ploidy values for the diploid and hypodiploid subclones. The outcomes from rcCAE and CHISEL deviated significantly from the anticipated results.

**Figure 4:**
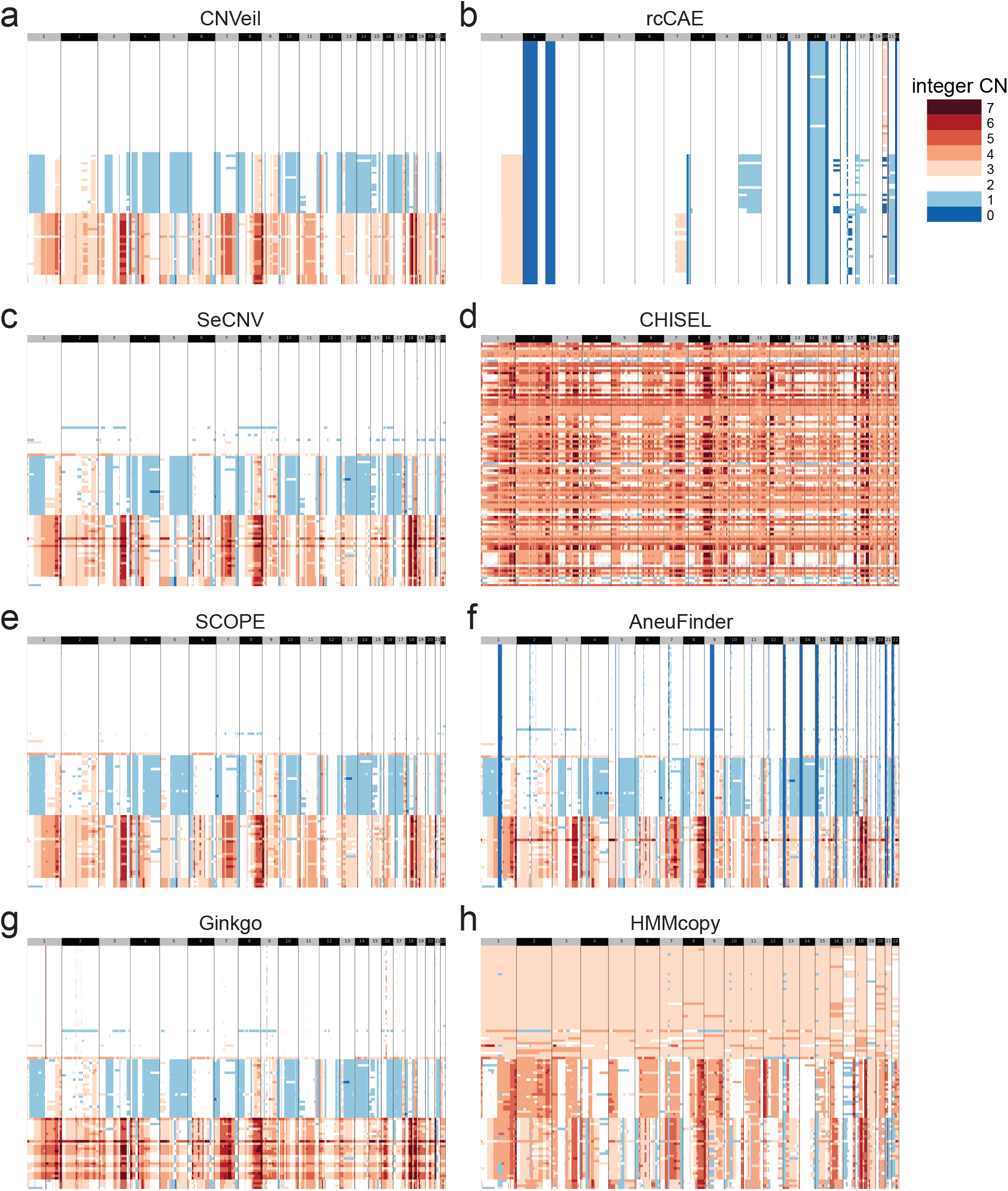
Comparison of inferred copy number profiles of single cells from the breast cancer patient T10. **(a-h)** Inferred copy number profiles by CNVeil, rcCAE, SeCNV, CHISEL, SCOPE, AneuFinder, Ginkgo, and HMMcopy. We ordered all heatmaps in a consistent cell order based on the order of reported subclones in FACS: normal cell subclone, one subclone of hypodiploid cancer cells, and two subclones of hyperdiploid cancer cells. Tools are shown in chronological order by publication year.

To further evaluate the performance of all tools quantitatively, we adopted CNV calls from array comparative genomic hybridization (aCGH) of purified bulk samples from the same patient T10 [30] as the gold standard. The aCGH data provided relative copy number information for each bulk sample. For each FACS-identified subclone, a corresponding gold standard CNV profile was established based on the product of the subclone’s ploidy and the relative copy number from aCGH data. By comparing estimated CNV profiles by each tool with the gold standard and using mean squared error (MSE) as the evaluation metric Figure (5), CNVeil demonstrated the lowest MSE among all eight tools across the four subclones.

**Figure 5:**
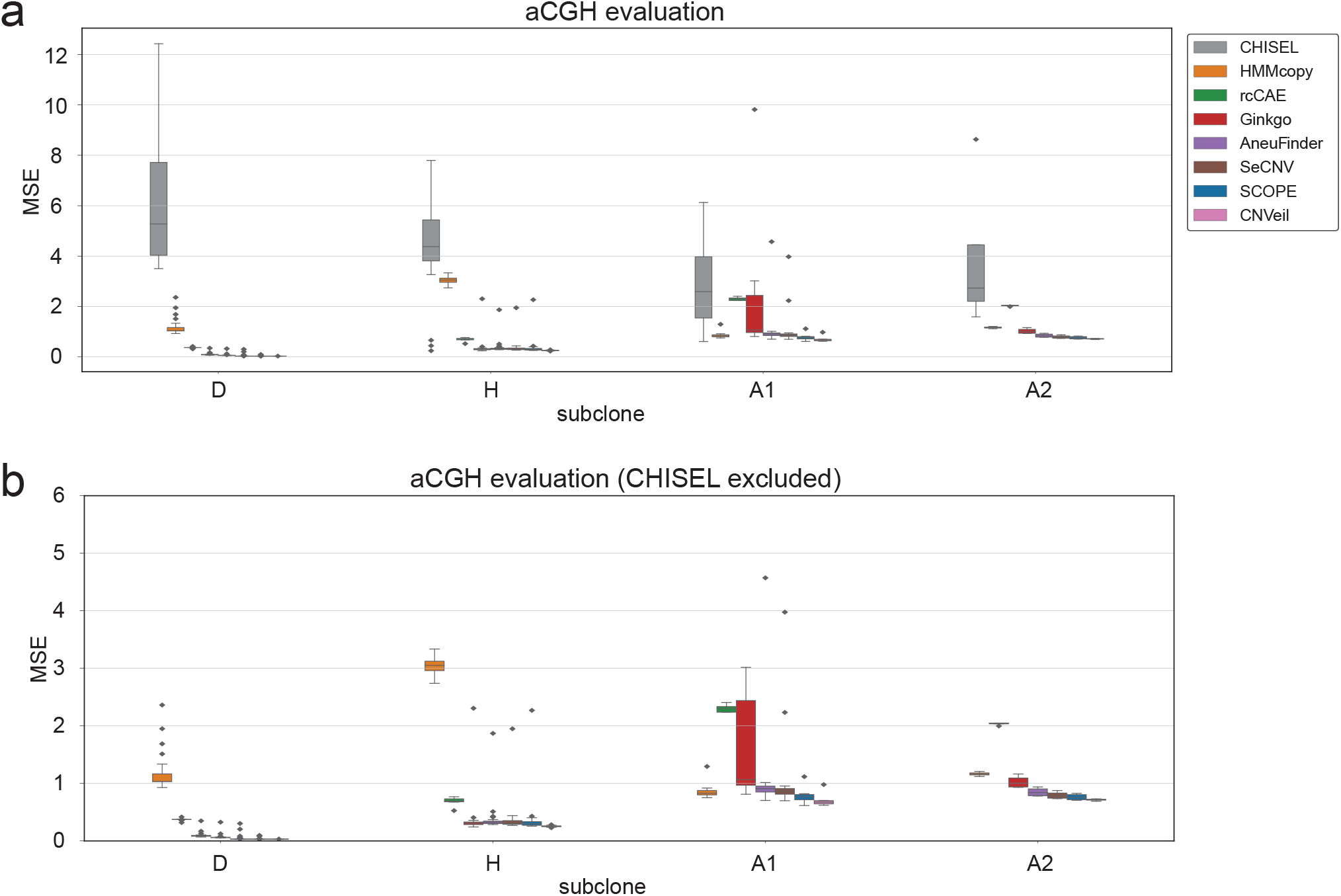
Orthogonal validation of single-cell copy number profiles by aCGH of purified bulk samples by FACS from T10). **(a)** Mean square error (MSE) plots for eight tools. **(b)** MSE plots for seven tools and CHISEL is excluded. The tools are ordered by average MSE in descending order.

### Evaluation on the scDNA-seq data of breast cancer patients T16

We also extended the analysis to another breast cancer patient, identified as T16 [31]. This dataset comprises 52 cells from primary tumor sites and 48 cells from metastatic tumor sites (Table 1). Both sites include a mixture of tumor and normal cells. Compared to T10, T16 demonstrated more homogeneous tumor subclones in both primary tumor and metastasis, indicating a monogenomic tumor.

The heatmaps in Figure 6 reveal that most tools demonstrated similar characteristics as what we observed in T10. Overall, CNVeil, SeCNV, SCOPE, AneuFinder, and Ginkgo showed a generally similar CNV profile pattern, distinguishing between a normal cell subclone and one subclone of hyperdiploid cancer cells. However, SeCNV, SCOPE, AneuFinder, and Ginkgo exhibited a few cells with incorrect ploidy and outlier bins across the genome. Upon careful inspection, CNVeil, SeCNV, SCOPE, AneuFinder, and Ginkgo all successfully identified two tumor subclones: primary and metastasis subclones (trisomic state). However, there were some disagreements regarding the copy number, especially on chromosomes 4, 5, and 7. CNVeil, SeCNV, AneuFinder, and Ginkgo predominantly predicted a copy number of 3 or 4 for most bins on chromosomes 4 and 5, whereas SCOPE predicted a copy number of 2. Similarly, for chromosome 7, CNVeil, SeCNV, AneuFinder, and Ginkgo predicted a copy number of 7, whereas SCOPE predicted a copy number of 5 or 6. In general, SCOPE tended to underestimate the copy numbers compared to CNVeil, SeCNV, and AneuFinder on T16 tumor cells. Conversely, the heatmap by HMMcopy showed a separation between normal and tumor cells, but it overestimated the ploidy of the normal cells to be 3. CHISEL showed a chaotic CNV profile, where there was no clear separation between normal and tumor cells. rcCAE failed to separate normal and tumor cells and underestimated the ploidy across all cells. In summary, CNVeil still demonstrated the best CNV profiles for subclone identification and segmentation.

**Figure 6:**
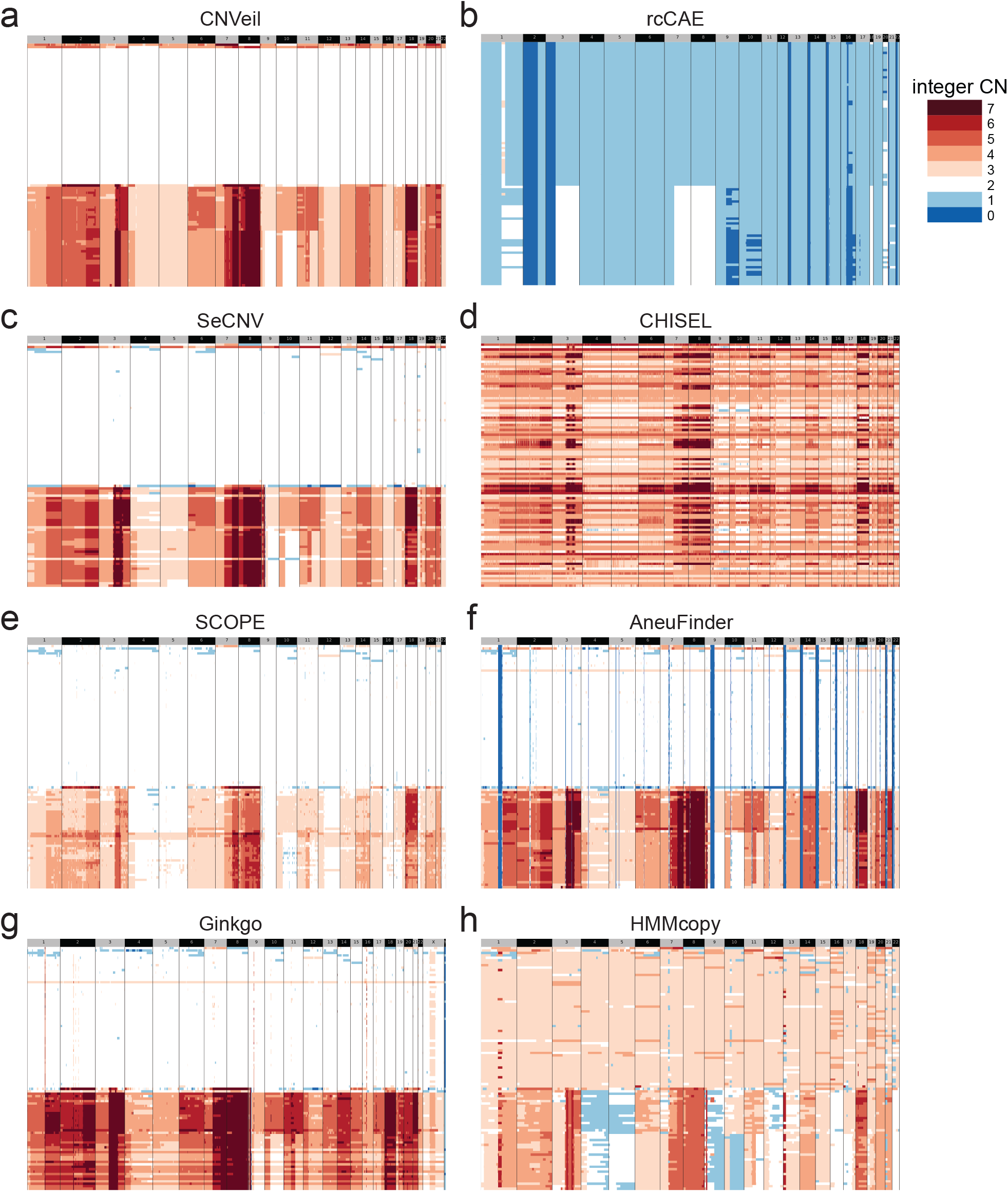
Comparison of inferred copy number profiles of single cells from the breast cancer patient T16. **(a-h)** Inferred copy number profiles by CNVeil, rcCAE, SeCNV, CHISEL, SCOPE, AneuFinder, Ginkgo, and HMMcopy. We ordered all heatmaps in a consistent cell order based on the hierarchical clustering results of the CNV profile by SCOPE. Tools are shown in chronological order by publication year.

### Evaluation on the scDNA-seq data of triple-negative cancer patient KTN302

We further applied the analysis to the scDNA-seq data of a triple-negative breast cancer patient, KTN302, whose treatment stages were well-documented [32]. This dataset includes 47 cells from the pre-treatment stage and 45 cells from the mid-treatment stage (Table 1).

To again ensure a fair and transparent comparison, we systematically ordered and organized cells in heatmaps based on the pre-treatment and midtreatment stages (Figure 7). We observed that CNVeil successfully detected a normal cell population from the mid-treatment stage and two subclones of aneuploid cells in the pre-treatment tumors. Although SCOPE showed a chaotic CNV profile at the mid-treatment stage, it was able to differentiate the tumor subclone from the normal cell population. Nevertheless, it is crucial to note that the KTN302 data is highly noisy. Therefore, SCOPE had to utilize mid-stage information as prior knowledge to select normal cells, achieving a reasonable outcome. When stage information was not provided to SCOPE, its CNV profile in the mid-treatment stage revealed many incorrect hyperdiploid cells (Figure S4). SeCNV, AneuFinder, and Ginkgo all exhibited numerous inaccuracies, including both hyperdiploid and hypodiploid cells during the mid-treatment stage, along with a few hypodiploid cells in the pre-treatment stage. These results indicated that when the scDNA-seq data was highly noisy, existing tools often failed to identify normal cells and correct bias, or they needed prior knowledge to assist these steps. CNVeil addressed this challenge by employing a novel PCA-based Gini coefficient to identify normal cells for bias correction. Moreover, CNVeil performed an initial normal-tumor cell clustering based on highly variable bins for robust ploidy estimation. CNVeil accurately detected 9 normal cells by PCA-based Gini coefficient and 43 out of 44 normal cells through the initial clustering (Table S1). This underscores CNVeil’s robustness and adaptability in various analytical scenarios. HMMcopy tended to overestimate the ploidy for the majority of cells in the mid-treatment stages. CHISEL showed an excessively noisy heatmap, while rcCAE exhibited an overly simplistic and smooth CNV profile.

**Figure 7:**
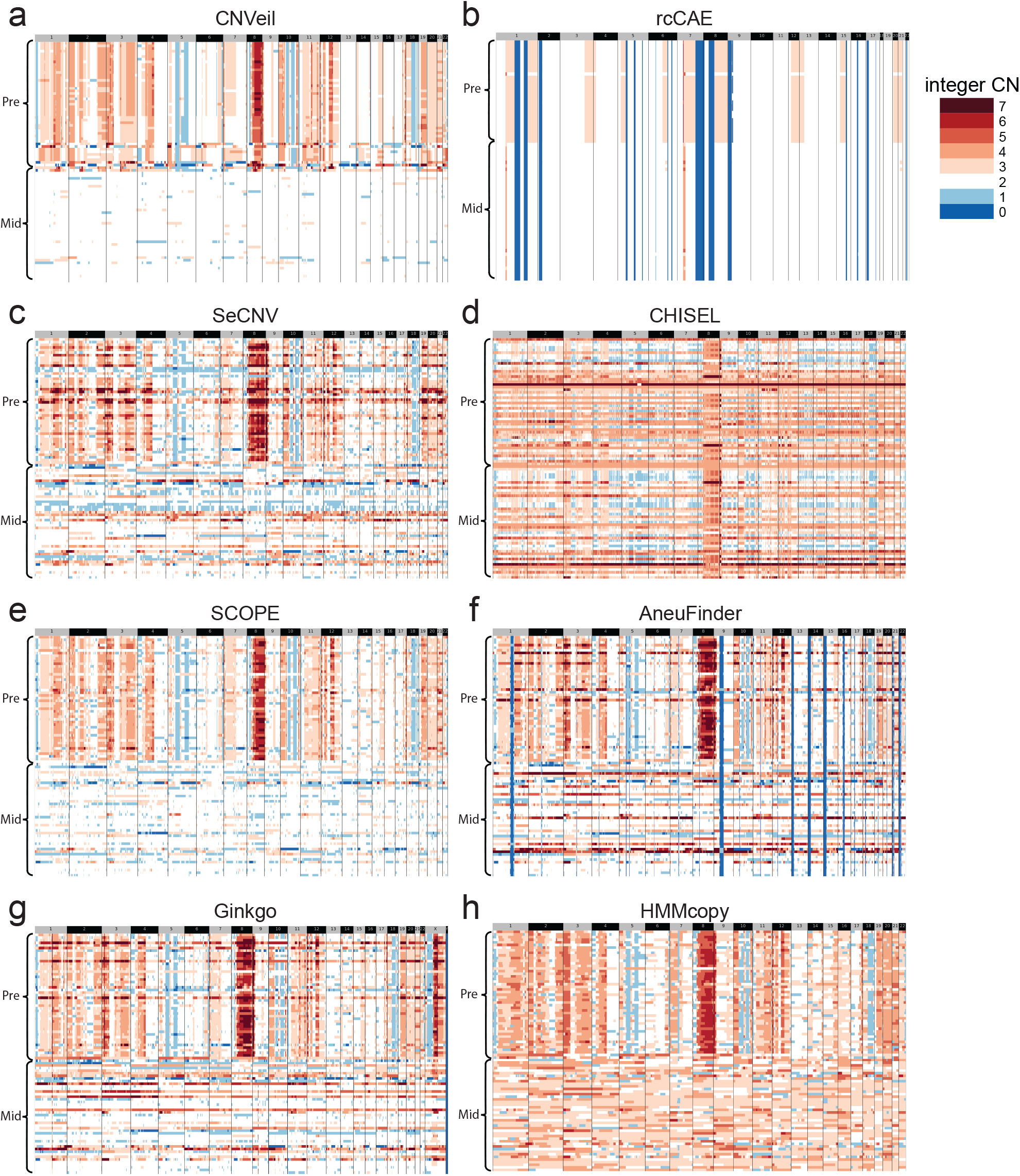
Comparison of inferred copy number profiles of single cells from a triple negative breast cancer patient KTN302. **(a-h)** Inferred copy number profiles by CNVeil, rcCAE, SeCNV, CHISEL, SCOPE, AneuFinder, Ginkgo, and HMMcopy. We ordered all heatmaps in a consistent cell order based on the pre-treatment and mid-treatment stages. Tools are shown in chronological order by publication year.

### CPU time and memory analysis

To provide users with an understanding of the efficiency of each tool in terms of CPU time and memory consumption, we recorded CPU time and memory usage across all benchmarked datasets (Figure 8 and Table S2). All tools utilized the single-cell BAM files as input and the CPU time was measured until each tool produced the final CNV profile. The memory usage was the peak memory during the whole procedure. Ginkgo is a web-based application and was excluded from this analysis.

**Figure 8:**
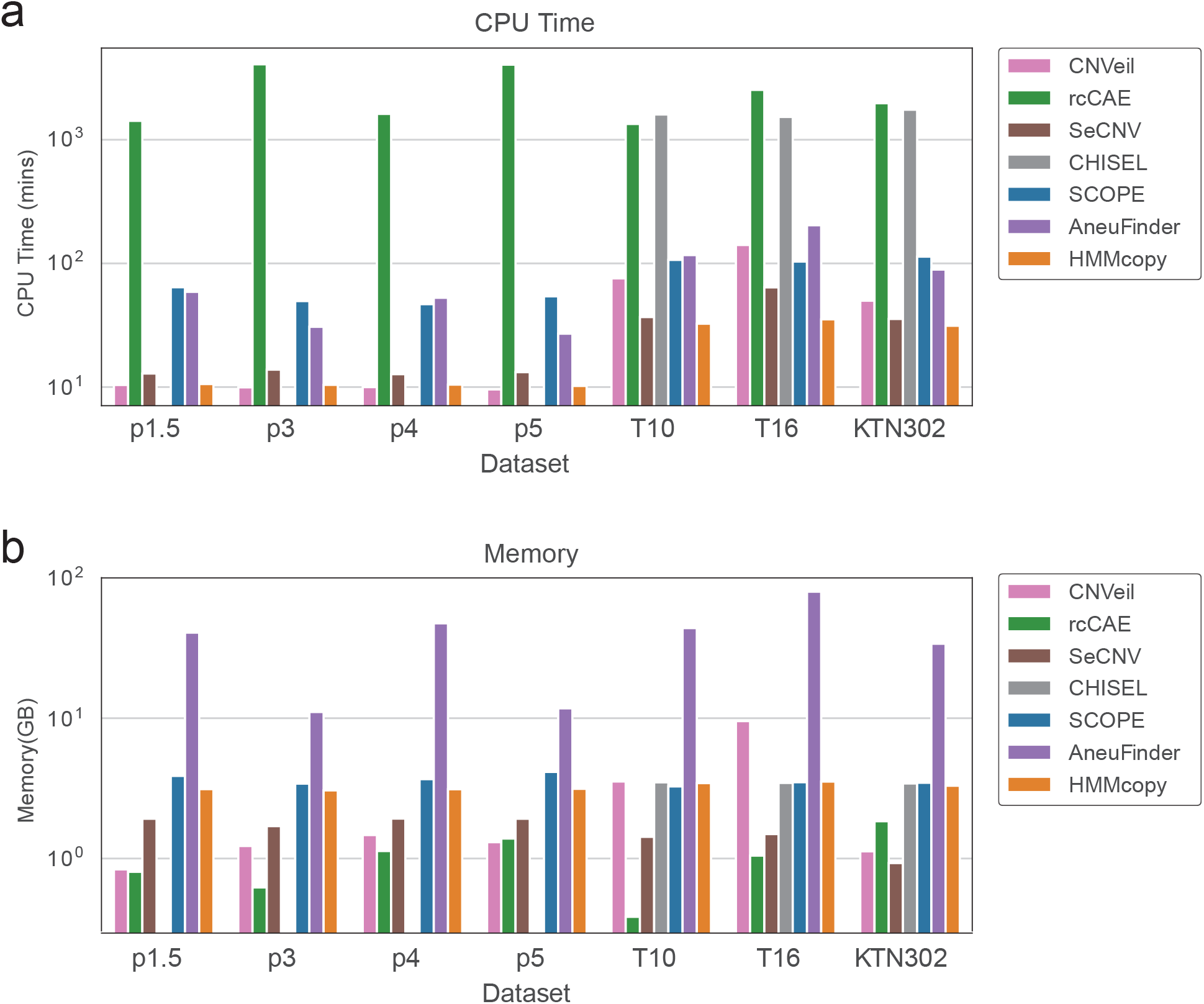
Comparison of different tools’ CPU time and memory usage across datasets. **(a)** CPU time in minutes (Mins) for eight tools across seven datasets. **(b)** Memory cost in gigabytes (GB) for eight tools across seven datasets. Tools are shown in chronological order by publication year.

CNVeil consumed CPU time within the range of 50 to 140 minutes, while maintaining a memory usage of less than 10 GB memory for real datasets. In terms of simulated datasets, CNVeil demonstrated efficient performance, requiring approximately 10 minutes of CPU time and utilizing less than 2 GB of memory. Overall, CNVeil ranked second or third places in terms of CPU time and memory usage across all datasets among all tools.

Furthermore, these CPU time and memory usage were measured while running CNVeil with 5 threads. In real practice, users with additional computational resources can enhance run-time efficiency by increasing the number of threads number. For example, when employing 20 threads, CNVeil completed tasks within 8 minutes for real data and 1 minute for simulation data, with a peak memory usage under 45 GB.

## Discussion

Here, we introduce CNVeil, a robust quantitative algorithm designed to accurately reveal CNV profiles from scDNA-seq data. To enhance the effectiveness of bias correction and address the challenges of the high noise in individual cells, CNVeil employs an innovative PCA-based Gini coefficient analysis to select normal cells. Variables such as subclonal ploidy, cell-specific ploidy, and the number of normal cells in the tumor sample determine the nature of the sample. Existing tools face challenges in accurately inferring ploidy for specific cells, largely due to sample variability induced by the complex nature inherent in cancer data. This often results in either underestimation or overestimation of copy numbers in certain cells. To address these challenges, CNVeil performs a multi-level hierarchical clustering, focusing on selected highly variable bins, to identify the normal subclone and different tumor subclones for robust ploidy estimation. For each cell, disregarding the variances of read depth in each bin, the product of cell-specific ploidy and normalized read depth yields the absolute copy number. However, existing methods for copy number inference are significantly impacted by the influence of variance and normalized read depth. To tackle this challenge, CNVeil incorporates a cross-cell consensus segmentation-based standardization to further normalize cell-specific read depth by reducing variance. Specifically, to identify the consensus segmentation, CNVeil refines subclones and introduces a novel change rate-based across-cell break-point identification approach to mitigate the effects of low coverage and data variability on a per-cell basis.

## Methods

### CNVeil workflow

The overall workflow of CNVeil consists of six interconnected and conceptual modules: a) Read count matrix construction; b) Data normalization by noise reduction and bias correction; c) Initial normal-tumor cell classification; e) Tumor subclones identification and ploidy estimation; f) Fine clustering and across-cell break-point and segmentation identification; h) Infer final CNV states. These are described in the following sections.

#### Read count matrix construction and removal of outlier bins and cells

Taking single-cell DNA sequencing data as input, CNVeil first aligns reads to the reference genome hg38 with BWA [33]. Those reads that cannot be uniquely aligned are removed. The genome is then divided into user-defined consecutive bins (500kb by default) and reads within a bin are aggregated (counted) to mitigate the impact of variable amplification and sequence sampling. The output is a cell-bin read count matrix. We denote the raw read count matrix as *RD ∈* ℝ^*n×m*^, where *n* is the number of cells and *m* is the number of genomic bins.

CNVeil drops the outlier bins with extreme GC contents (*<*20% or *>*80%), and the outlier bins with mappability below 90%. Cells with mapped reads proportion (the ratio of the number of reads mapped to reference to the number of all reads) less than a threshold (0.3 by default) are also removed as outliers for quality control.

#### Data normalization by noise reduction and bias correction

In single-cell cancer genomics studies, diploid cells often serve as normal controls for read depth normalization [19,27]. Nevertheless, information regarding case-control labeling and cell ploidy might not always be readily available. Cell-specific Gini coefficients to quantify the variability within individual cells can be used to select normal cells out of the entire cell population by empirical evidence [19]. However, noise within read depth signals presents a significant challenge in this analysis [4]. To overcome this, CNVeil employs a Principal Component Analysis (PCA) based Gini coefficients to discern the substantive signal from the pervasive noise, thereby identifying bins marked by notable variance to compute Gini coefficients. To achieve this, CNVeil first performs PCA [34] on the read count matrix *RD*, generating a matrix of principal components. CNVeil focuses on the first component, PC[:, 1], which carries the largest data fluctuation information. CNVeil then selects the bins that have higher weights along the first component. Consider the read depth vector for a specific cell *i* and a weight vector **w**, the weighted sum for selected bins based on high PCA scores can be denoted as:

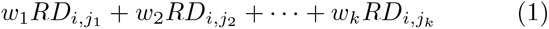

where *j*_1_, *j*_2_, …, *j*_*k*_ are the indices of bins chosen by PCA, *{j*_1_, *j*_2_, …, *j*_*k*_*} ⊆ {*1, 2, 3, …, *M}*. Each weight *w*_*j*_ corresponds to the importance of the *j*-th bin in the PCA score ranking. Bins contribute more to the first component, to be specific, the first 40% will be saved for the analysis.

The PCA-based Gini coefficient for each cell *i* is computed as follows:

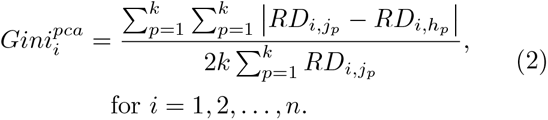

In this context, cells exhibiting a PCA-based Gini coefficient below 0.12 are classified as normal cells. In several real data, we observed PCA-based Gini coefficient served as a more effective metric to differentiate normal cells than Gini coefficient. In practice, CNVeil does not require the identification of all diploid cells from the cell population, only requiring a subset of all normal cells to serve as normal controls. After obtaining normal cells, CNVeil corrects for the bias of each bin and generates the normalized read count matrix.

This process normalizes the read depth of each cell by a bias scale from normal controls, thus achieving uniformity across the data. The bias scale can be computed by the equation:

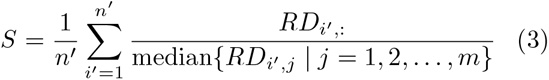

where 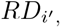 : is a vector representing read depth across all bins for normal cell *i*^*′*^, and *n*^*′*^ is the number of normal cells. *S* is the bias scale vector, and *S*_*j*_ is the bias scale for bin *j*. To normalize read depth by bin-wise, the operation can be mathematically represented as follows:

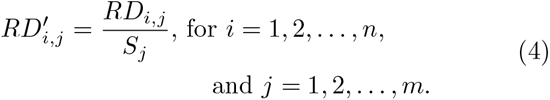

where *RD*^*′*^ is the matrix of normalized read depths.

This scale captures the typical deviation patterns inherent in normal cells. Deploying this bias scale across the data adjusts for systemic biases. At the end of this module, CNVeil generates a normalized read depth matrix *RD*^*′*^ that is noise-reduced and bias-corrected. This matrix serves as the input for the subsequent modules.

#### Hierachical clustering for different levels of cell clusters

CNVeil employs the agglomerative clustering algorithm [35] to perform different levels of cell clustering including normal-tumor cell clustering, tumor cell subclone identification, and fine clustering within subclones. For each cell, if we disregard the variances of read depth in each bin, the absolute copy number is the product of cell-specific ploidy and normalized read depth. Thus, to infer CNV states, we aim to perform different levels of clustering to either estimate optimal ploidy or conduct cross-cell breakpoint and segmentation identification to further standardize read depth.

The main idea behind agglomerative clustering is to initialize each cell as an individual cluster and then iteratively merge clusters based on their similarity measure until only one big cluster containing all the cells remains in the final stage. This process creates a tree-like structure, commonly known as a dendrogram, which can be cut at different heights to obtain different clustering results. The height of a certain level in a dendrogram represents the linkage distance of clusters at that level. Intuitively, cutting the dendrogram at a lower height generates a more granular clustering result, while at a higher height generates a more coarser clustering result. Ward’s linkage distance is used in CNVeil pipeline to measure the linkage distance. It can be calculated as

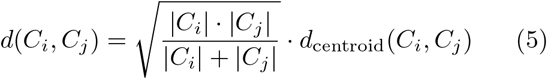

Where d(*C*_*i*_,*C*_*j*_) is the Ward’s linkage distance between clusters *C*_*i*_ and *C*_*j*_, |*C*_*i*_| and |*C*_*j*_| are the sizes of clusters *C*_*i*_ and *C*_*j*_ which are defined by the number of cells within each cluster, and *d*_*centroid*_(*C*_*i*_,*C*_*j*_) is the Euclidean distance between the centroids of clusters *C*_*i*_ and *C*_*j*_.

#### Initial normal-tumor cell classfication

To perform the agglomerative algorithm for the initial normal-tumor cell clustering, CNVeil specifically selects the highly variable bins (across cells) as the feature vector to represent each cell, instead of using all consecutive bins. Highly variable bins exhibit significant variability in their read depth levels across cells.

To select highly variable bins, CNVeil takes the normalized read depth matrix *RD*^*′*^ as input and computes the variance across cells for each bin, thereby identifying those bins that exhibit significant variability. The variance for a given bin *j* can be expressed as:

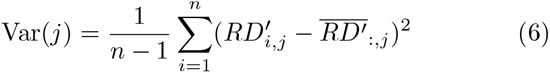

where 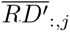 is the average read depth for bin *j, i* is the cell index, and *n* is the number of cells. CNVeil identifies top 10% of these bins with the highest variance as highly variable bins. These bins are of particular interest because they often contribute to the biological diversity and heterogeneity observed among individual cells. Highly variable bins can serve as potential marker bins for specific cell types. The normalized read depth matrix composed of only highly variable bins is denoted as 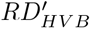. CNVeil, therefore, taking 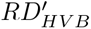 as input, uses agglomerative clustering to separate all cells into two clusters at the initial stage by setting the number of clusters equal to 2.

Next, CNVeil utilizes a numerical optimization method to determine the optimal ploidy level for each cluster. The cluster that is determined to have ploidy level close to 2 will be regarded as normal cell cluster, while the other will be treated as a tumor cell cluster. Specifically, CNVeil picks the cluster-level ploidy that minimizes the squared error that is quantified between the predicted CNVs and their nearest integer values across all bins. Identifying the optimal ploidy that results in the least discrepancy allows CNVeil to accurately infer the genomic state of the cell cluster under examination. The numerical optimization procedure is described below.

CNVeil defines a candidate ploidy set *Ploidy* = [1.50, 1.55, …, 5.50], which ranges from 1.50 to 5.50 with a step size of 0.05. |*P* | = *K*. The Scaled Copy Number Profile (SCNP) is given by:

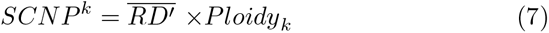

where 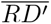 is a vector that represents the average read depth across all cells for each bin. The Sum of Squares (SoS) is calculated as:

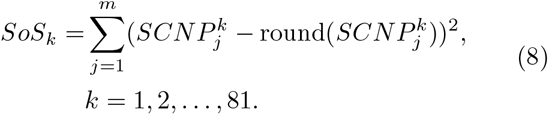

The optimal ploidy level for each cluster, represented by *ploidy*^*∗*^ or *Ploidy*[*k*], can be achieved by the value of *k* that minimizes the Sum of Squares (SoS):

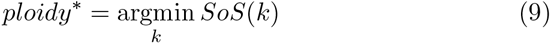

At this module, CNVeil performs agglomerative clustering relying on highly variable bins to differentiate cells into two clusters and further utilizes the numerical optimization method to determine the optimal ploidy level for each cluster to classify normal and tumor cell clusters.

#### Tumor subclones identification and ploidy estimation

In practice, a patient’s tumor cells may contain multiple subclones, which could either share similar ploidy levels or have distinct ploidy levels [36–38]. Therefore, it is important to identify tumor subclones and estimate each subclone’s ploidy level reliably. However, ploidy estimation based on a small number of cells (for example, less than 5) may lack accuracy due to the inherent noise present in single-cell sequencing data. It is more likely to recover the actual ploidy level when a larger amount of cells that share the same ploidy level are provided [39]. Hence, in this step, when performing hierarchical clustering, we try to maximize the size of each cluster under the constraint that cells in one cluster share the same ploidy level. Quantitatively, an estimation of the minimal Ward’s distance between any two tumor subclones that have different ploidy values was empirically drawn from the experiments of multiple real datasets (T10, T16, KTN302, etc). This minimal distance is denoted as *D*_*p*_ and is estimated to be around 30. The choice of *D*_*p*_ is affected by the number of highly variable bins. Given the distance constraint *D*_*p*_, the agglomerative clustering is applied to the tumor cells subset of the matrix 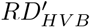, generating multiple ploidy-consistent tumor subclones. Subsequently, the aforementioned numerical optimization method is applied to each subclone to estimate the optimal ploidy *ploidy*^*∗*^, which will be used to infer CNV states in the last module.

#### Fine clustering and across-cell breakpoint and segmentation identification

Cells within the same sub-clone share the same cellular breakpoints and a common evolutionary history [6, 40]. To enhance segmentation accuracy by addressing the challenges posed by investigating each cell independently, CNVeil introduces a cross-cell consensus segmentation approach for identifying breakpoints and segmentation across the genome. To maximize cell-level variation for break-points, CNVeil further performs fine clustering on each tumor subclone before performing cross-sample break-point detection.

Similar to the preceding round of coarse clustering to identify tumor subclones, an empirical threshold *D*_*b*_ is determined. This threshold represents the minimal Ward’s distance between any two cell clusters that have different breakpoint profiles (where CNV state changes) and is estimated to be approximately 14. Given *D*_*b*_, another round of agglomerative clustering is applied to the tumor subset of the matrix 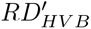 to generate more refined clusters. At the end of this step, each generated cluster is expected to share a common breakpoint profile.

Next, CNVeil performs cross-cell breakpoint detection within each cell cluster. This algorithm assesses the statistical significance of change points by aggregating read depth across cells within two target windows around each point and determining the significant changes between target windows across all points. For each cell cluster, CNVeil takes the normalized read depth matrix as the input, represented by a *p × m* matrix (*p* is the number of cells in each cluster and *m* is the number of genomic bins). Given *m* bins, (*m −* 1) boundary points of all bins could be potential change points. We denote these (*m−*1) points as a set *S*_*b*_. CN-Veil then uses a sliding window of size *p × w* (*w* is the window size, and it is set as 6 bins by default) to investigate the read depth changes around those potential change points. Specifically, for the boundary point between *bin*_*k*_ and *bin*_*k*+1_, CNVeil calculates the median read depth in the left window (from *bin*_*max*(*k−w*+1,1)_ to *bin*_*k*_) as *MRD*_*left*_ and the median read depth from the right window (from *bin*_*k*+1_ to *bin*_*min*(*k*+1+*w,m*)_) as *MRD*_*right*_.

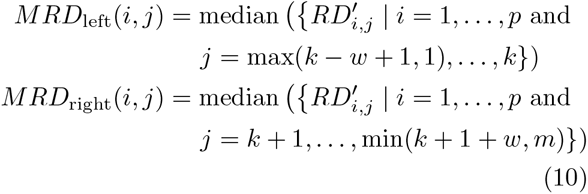

Next, CNVeil calculates the read depth change rate as below:

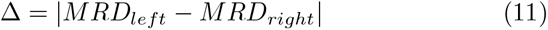

CNVeil collects Δ for each boundary point in every cell cluster, generating a set of read depth change rates denoted as *S*_Δ_. The changing rate threshold *t*_Δ_ is established as the 90% quantile of *S*_Δ_. Any boundary point in *S*_*b*_ where the changing rate exceeds the threshold *t*_Δ_ is identified and collected as a change point.

Within each cluster, a chaining algorithm is then applied to chain the change points into multiple disjoint regions. The chaining algorithm establishes an edge between any two change points if their distance is equal to one bin size, generating a network of multiple disjoint components. Each component forms a region that encompasses a real breakpoint in our assumption.

To control the false discovery rate of breakpoints, regions that contain less than six change points are excluded from the subsequent analysis since they are likely the artifact of sequencing noise. Finally, CNVeil selects the middle change point in each region as the predicted breakpoint. CNVeil repeats the cross-cell breakpoint detection procedure for each cell cluster and generates a unique breakpoint profile for each cluster.

Finally, the segmentation is established by defining a segment as the consecutive bins between every two predicted breakpoints. CNVeil also performs fine clustering and across-cell breakpoints and segmentation identification for the initial normal cell cluster with the same approach.

#### Infer final CNV states

In the final module of the pipeline, the objective of CNVeil is to infer the copy number state for each bin of each cell. As we discussed before, for each cell, if we disregard the variances of read depth in each bin, the product of cell-specific ploidy and normalized read depth is the absolute copy number. To further normalize the read depth and minimize variance, CNVeil utilizes the cross-cell segmentation achieved in the previous module to standardize each normalized read depth 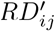 as below:

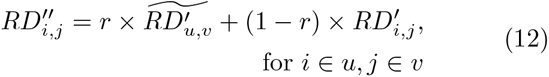

where 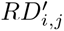 is the normalized read depth for the *i -* th cell and *j*-th bin, and 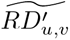 is the median read depth for the *v*-th segment in the *u*-th fine cell cluster. 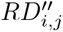 is the standardized read depth, and *r* is the standardization factor (0.5 by default). By standardizing, CNVeil ensures 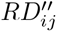 faithfully represents genuine genomic aberrations while minimizing the impact of artifact noise.

The Final Copy Number Profile (FCNP) is then inferred as:

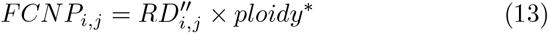

CNVeil uses subclone ploidy^*∗*^ to represent cell-specific ploidy because 1) ploidy^*∗*^ is the optimal ploidy when maximizing the size of each cluster under the constraint that cells in one cluster share the same ploidy level; 2) although there is variability in read depth across all bins for each cell within the subclone, the cell-specific ploidy does not exhibit significant variability.

#### Simulated data by simSCSnTree

We employed Sim-SCSnTree to generate simulated datasets with a gold standard. The simulation process executed by SimSCSnTree is intricately designed in three primary steps, reflecting the complexities inherent in cancer genomics.

#### Step 1: Phylogenetic tree construction and genomic variation modeling

In the initial step, a phylogenetic tree is crafted using SimSCSnTree, with each node representing a distinct genome segment, and the connecting edges denoting genetic variations (CNVs). Built upon the hg19 reference genome, it ensures our simulation mirrors actual genomic structures and sequences. The outcome produces two critical numpy (.npy) files: one delineating the CNVs alongside the tree structure and another storing intermediate information essential for the subsequent simulation step.

SimSCSnTree controls the characteristics of the tree and the final CNV profile by multiple parameters, such as the multiplier of the mean CNV on the root, the rate of deletion, the mean, and median of the tree width distribution, etc. For example, to generate the tree for a dataset with an average ploidy of 1.5, we used the command below:

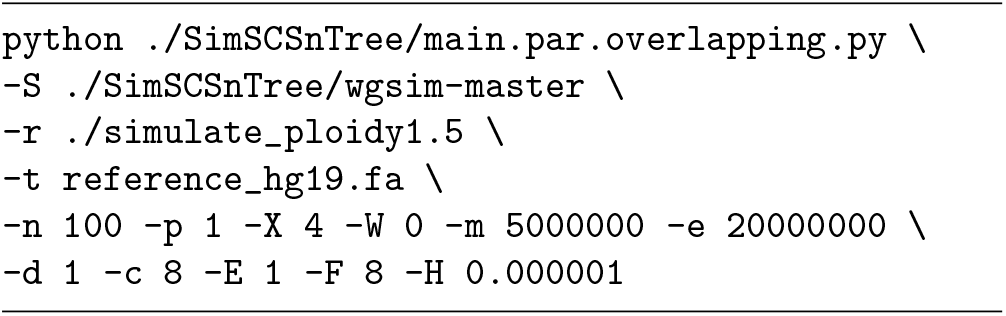

where **-n** is number of cells, **-p** is number of processors, **-X** is the multiplier of the mean CNV on root, **-W** denotes whether there is whole chromosome amplification (0 indicating no), **-m** is the minimum copy number size, **-e** is the parameter for the exponential distribution for copy number size beyond the minimum one, **-d** is the rate of deletion, **-c** is the average number of copy number variations to be added on a branch, **-E** is the whole amplification copy number addition, **-F** is the mean of the tree width distribution, and **-H** is the standard deviation of the tree width distribution.

To generate a dataset of other ploidy, the user needs to adjust the corresponding parameters. To reproduce all four simulated datasets in this paper, users can refer to our GitHub for more details.

#### Step 2: Read sampling from the simulated genome building upon the tree

In the second step, reads are sampled from the genome at selected nodes of the tree. This phase is critical in translating theoretical genomic information into practical sequencing data. By utilizing the .npy files generated from the first step, this phase simulated the sequencing process, closely resembling real-world sequencing techniques.

For example, to simulate reads from the tree generated by step1, we used the command below:

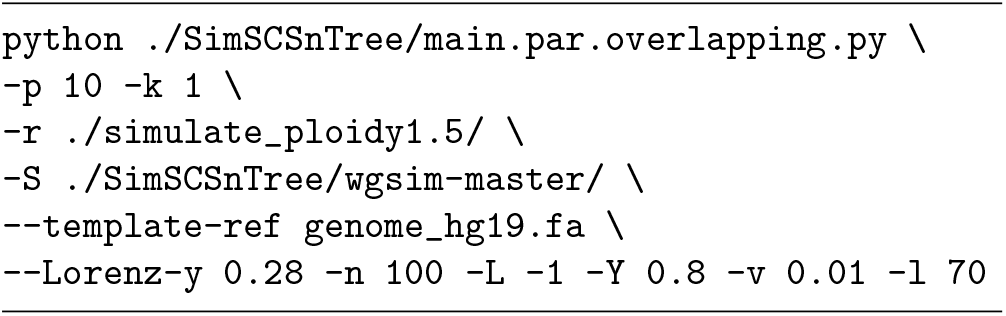

where **-p** is the number of processors, **-k** denotes whether to skip step 1 (1 for skip), **–Lorenz-y** is the value on the y-axis of the Lorenz curve, **-n** is number of cells, **-l** is the read length, **-v** is the average coverage of the sequence, **-L** controls the levels to be sequenced, and **-Y** specifies a range of nodes to sequence. the above command for step 2 was employed for generating all four simulated datasets in this paper.

#### Step 3: Extract the gold standard set

The ground truth data for CNVs obtained from this simulation served as the gold standard against which the accuracy and effectiveness of various CNV inference tools can be assessed. The ground truth was extracted from different nodes (tumor subclones), that contain a set of breakpoints (where the copy number changes) and the left and right copy numbers for each breakpoint. Similarly, the callset from each CNV inference tool was also reformatted into a set of breakpoints and neighboring copy numbers for a fair comparison and evaluation.

#### Evaluation against gold standard in simulated datasets

In simulated data, we compared the callset by each tool with the gold standard set using two evaluation modes: segmentation mode and CNV mode.

In segmentation evaluation mode, we required the position of the called breakpoint and gold standard breakpoint to approximately align. The called break-point matches the gold standard breakpoint if they satisfy the below conditions:

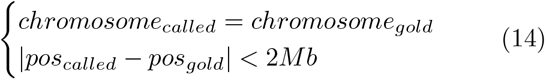

In stringent CNV evaluation mode, we required both the breakpoints and the copy number on the left and right bins of the breakpoint to match between the called event and the gold standard. The called event matches the gold standard if they satisfy the below conditions:

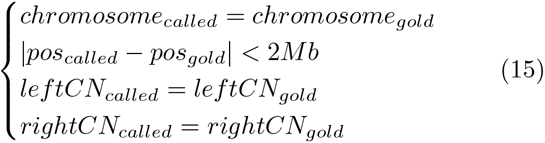

Following an N-to-N matching procedure between each callset and the gold standard set, breakpoints from the callset matched to any breakpoint in the gold standard set are classified as true positives (TPs), while the remaining breakpoints from the callset are considered false positives (FPs). Conversely, break-points from the gold standard set without a matching counterpart in the callset are classified as false negatives (FNs). The evaluation matrices are calculated as below:

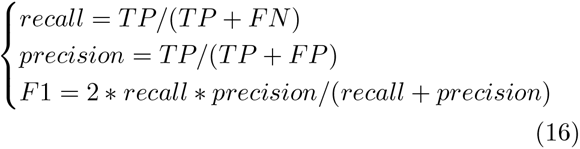

## Code availability

All code is available at https://github.com/maiziezhoulab/CNVeil. We implemented both python and R version of CNVeil for users. The computation cost was benchmarked in the Python version.

## Acknowledgements

This work was supported by the NIH NIGMS Maximizing Investigators’ Research Award (MIRA) R35 GM146960 to X.M.Z.

## Author’s contributions

X.M.Z. conceived and led this work. W.Y., C.L and X.M.Z. designed the framework and W.Y. and C.L implemented the framework. W.Y., C.L., Y.H., L.Z., Z.W., Y.H.L performed data analysis. L.Z. and X.M. assisted with the simulation analysis. W.Y., C.L, and X.M.Z. wrote the manuscript with input from all authors.

## Competing interests

The authors declare that they have no competing interests.

## Notes

### Competing Interest Statement

The authors have declared no competing interest.

