## Appendix for "CNVeil enables accurate and robust tumor subclone identification and copy number estimation from single-cell DNA sequencing data"

Yuan et al.

### 1 Supplementary Tables

| Datasets | Two procedures in CNVeil | Gold | True Positive | False Positive | False Negative | Recall | Precision | F1-score |
|---|---|---|---|---|---|---|---|---|
| T10 | PCA-based Gini | 45 | 41 | 0 | 4 | 91.11% | 100.00% | 95.35% |
| T10 | Initial Clustering | | 45 | 0 | 0 | 100.00% | 100.00% | 100.00% |
| KTN302 | PCA-based Gini | 44 | 9 | 0 | 35 | 20.45% | 100.00% | 33.96% |
| KTN302 | Initial Clustering | | 43 | 7 | 0 | 100.00% | 86.00% | 92.47% |
| sim p1.5 | PCA-based Gini | 50 | 50 | 0 | 0 | 100.00% | 100.00% | 100.00% |
| sim p1.5 | Initial Clustering | | 50 | 0 | 0 | 100.00% | 100.00% | 100.00% |
| sim p3 | PCA-based Gini | 50 | 50 | 0 | 0 | 100.00% | 100.00% | 100.00% |
| sim p3 | Initial Clustering | | 50 | 0 | 0 | 100.00% | 100.00% | 100.00% |
| sim p4 | PCA-based Gini | 50 | 50 | 0 | 0 | 100.00% | 100.00% | 100.00% |
| sim p4 | Initial Clustering | | 50 | 0 | 0 | 100.00% | 100.00% | 100.00% |
| sim p5 | PCA-based Gini | 50 | 50 | 0 | 0 | 100.00% | 100.00% | 100.00% |
| sim p5 | Initial Clustering | | 50 | 0 | 0 | 100.00% | 100.00% | 100.00% |

Table S1: **Performance metrics for normal cells identification by two procedures in CNVeil across different datasets.** For each dataset, this table summarizes several metrics. True Positive cells: cells correctly identified as normal by CNVeil; False Positive cells: cells incorrectly identified as normal by CNVeil; False Negative cells: normal cells not identified by CNVeil. “Gold”: actual number of normal cells in each dataset. Recall, precision, and F1 scores are also listed in the table.

| Tool | Dataset | CPU Time(Mins) | Memory(GB) |
|---|---|---|---|
| CNVeil | T10 | 75.68 | 3.54 |
| rcCAE | T10 | 1340.93 | 0.38 |
| SeCNV | T10 | 36.82 | 1.43 |
| CHISEL | T10 | 1599.98 | 3.46 |
| SCOPE | T10 | 106.73 | 3.26 |
| AneuFinder | T10 | 116.73 | 3.26 |
| HMMcopy | T10 | 32.53 | 3.45 |
| CNVeil | T16 | 140.85 | 9.54 |
| rcCAE | T16 | 2528.25 | 1.05 |
| SeCNV | T16 | 63.98 | 1.49 |
| CHISEL | T16 | 1528.85 | 3.49 |
| SCOPE | T16 | 103.45 | 3.50 |
| AneuFinder | T16 | 203.52 | 79.73 |
| HMMcopy | T16 | 35.28 | 3.54 |
| CNVeil | KTN302 | 49.93 | 1.13 |
| rcCAE | KTN302 | 1972.92 | 1.84 |
| SeCNV | KTN302 | 35.55 | 0.93 |
| CHISEL | KTN302 | 1748.98 | 3.42 |
| SCOPE | KTN302 | 113.40 | 3.47 |
| AneuFinder | KTN302 | 89.15 | 33.95 |
| HMMcopy | KTN302 | 31.32 | 3.30 |
| CNVeil | sim p1.5 | 10.40 | 0.84 |
| rcCAE | sim p1.5 | 1420.85 | 0.80 |
| SeCNV | sim p1.5 | 12.87 | 1.92 |
| SCOPE | sim p1.5 | 64.15 | 3.88 |
| AneuFinder | sim p1.5 | 58.91 | 40.72 |
| HMMcopy | sim p1.5 | 10.58 | 3.12 |
| CNVeil | sim p3 | 9.95 | 1.23 |
| rcCAE | sim p3 | 4071.77 | 0.62 |
| SeCNV | sim p3 | 13.84 | 1.70 |
| SCOPE | sim p3 | 49.60 | 3.42 |
| AneuFinder | sim p3 | 30.72 | 11.07 |
| HMMcopy | sim p3 | 10.42 | 3.06 |
| CNVeil | sim p4 | 9.99 | 1.47 |
| rcCAE | sim p4 | 1617.10 | 1.13 |
| SeCNV | sim p4 | 12.72 | 1.93 |
| SCOPE | sim p4 | 46.80 | 3.68 |
| AneuFinder | sim p4 | 52.73 | 47.46 |
| HMMcopy | sim p4 | 10.47 | 3.12 |
| CNVeil | sim p5 | 9.57 | 1.31 |
| rcCAE | sim p5 | 4044.58 | 1.39 |
| SeCNV | sim p5 | 13.20 | 1.92 |
| SCOPE | sim p5 | 54.17 | 4.14 |
| AneuFinder | sim p5 | 27.03 | 11.76 |
| HMMcopy | sim p5 | 10.22 | 3.14 |

Table S2: **Computing resource consumption of CNV inference tools.** This table lists the run time and memory usage of seven different CNV inference tools across three real and four simulated scDNA-seq datasets. The CPU time is reported in minutes (Mins), and the memory usage is provided in gigabytes (GB).

#### 2 Supplementary Figures

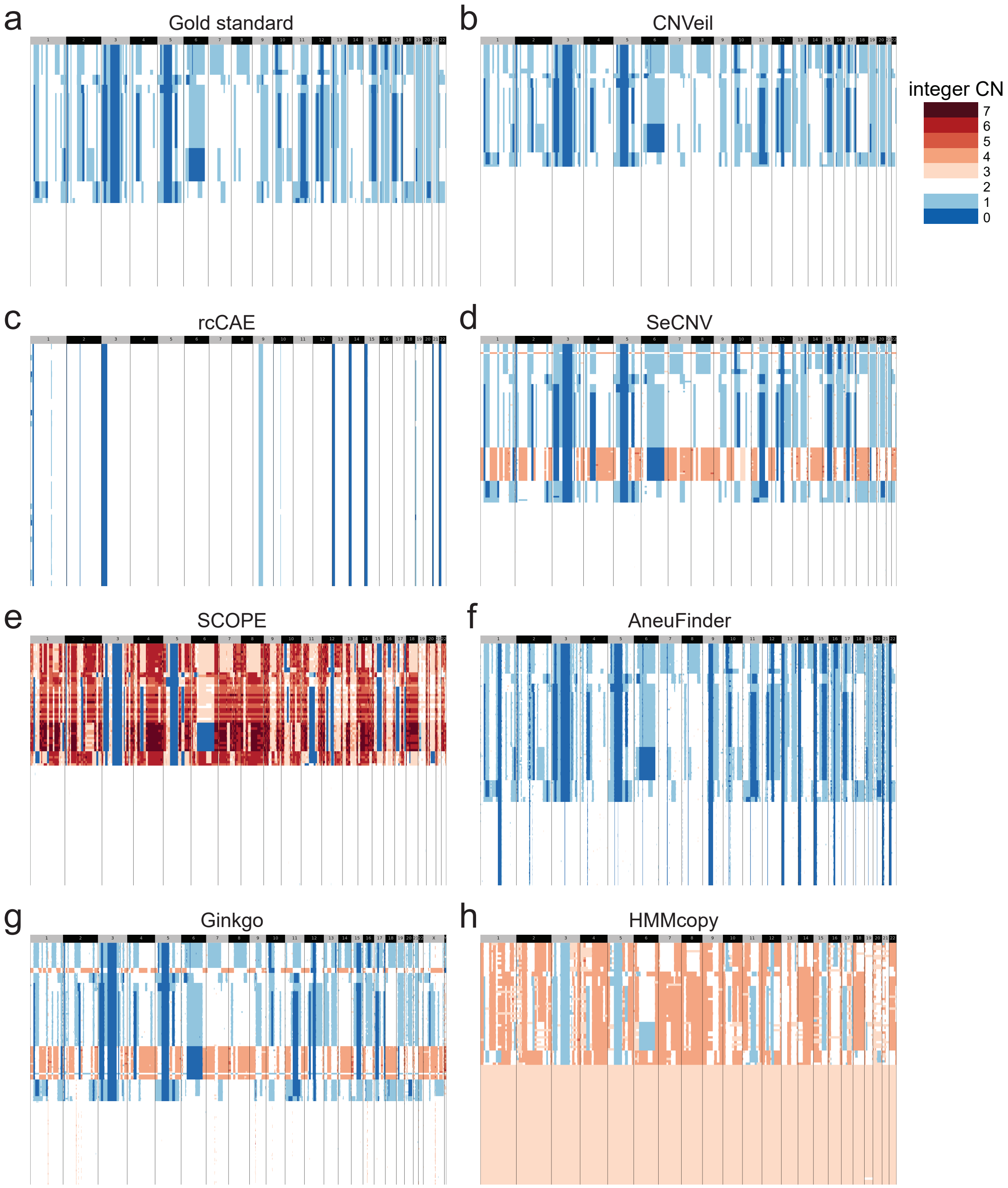

Figure S1: **Comparison of inferred copy number profiles of single cells from the simulated dataset with a ploidy of 1.5. (a)** Established copy number profiles by the gold standard. **(b-h)** Inferred copy number profiles by CNVeil, rcCAE, SeCNV, SCOPE, AneuFinder, Ginkgo, and HMMcopy. We ordered all heatmaps in a consistent cell order aligned with the gold standard which includes a normal cell subclone and a tumor subclone. Tools are shown in chronological order by publication year.

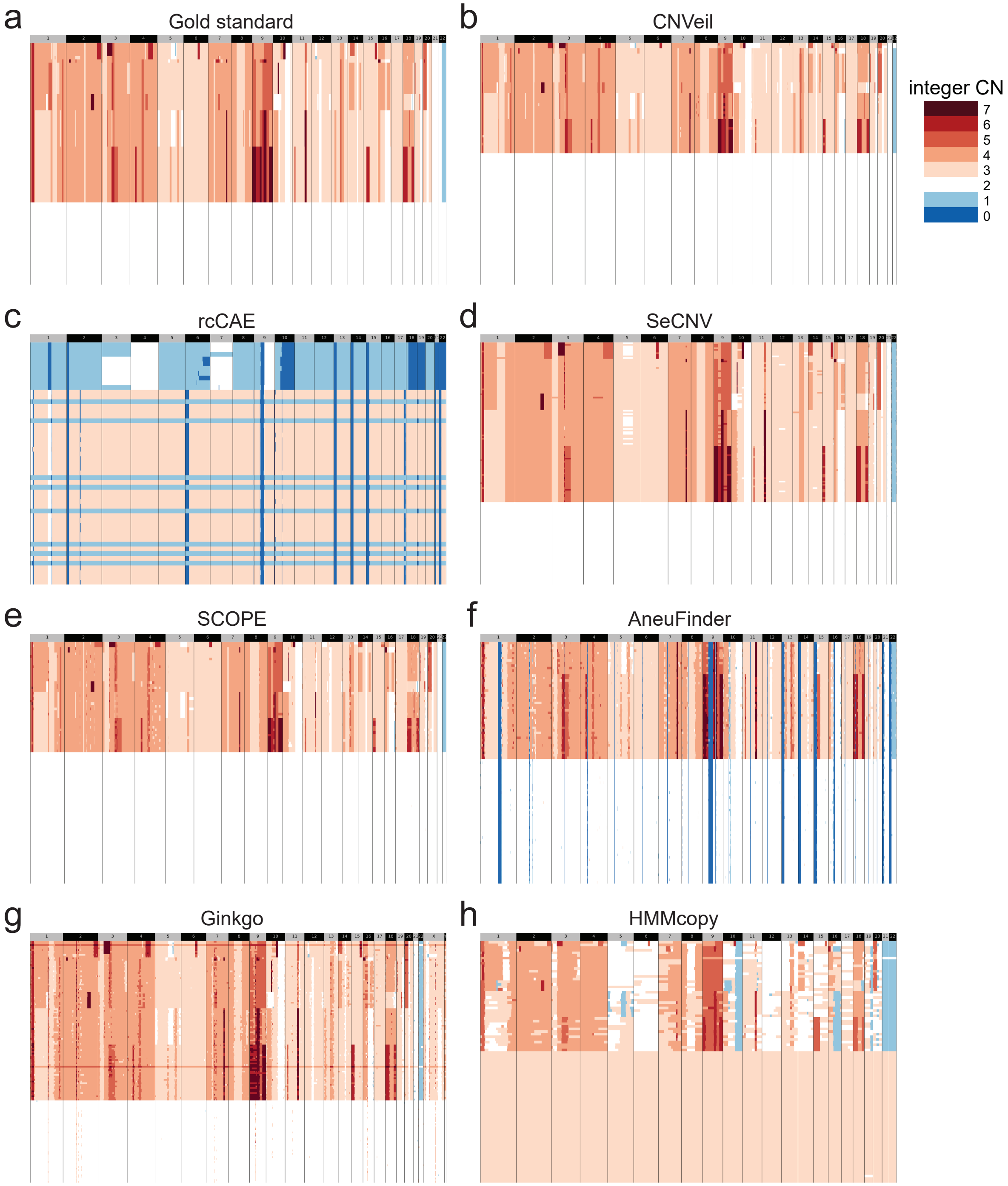

Figure S2: **Comparison of inferred copy number profiles of single cells from the simulated dataset with a ploidy of 3.0. (a)** Established copy number profiles by the gold standard. **(b-h)** Inferred copy number profiles by CNVeil, rcCAE, SeCNV, SCOPE, AneuFinder, Ginkgo, and HMMcopy. We ordered all heatmaps in a consistent cell order aligned with the gold standard which includes a normal cell subclone and a tumor subclone. Tools are shown in chronological order by publication year.

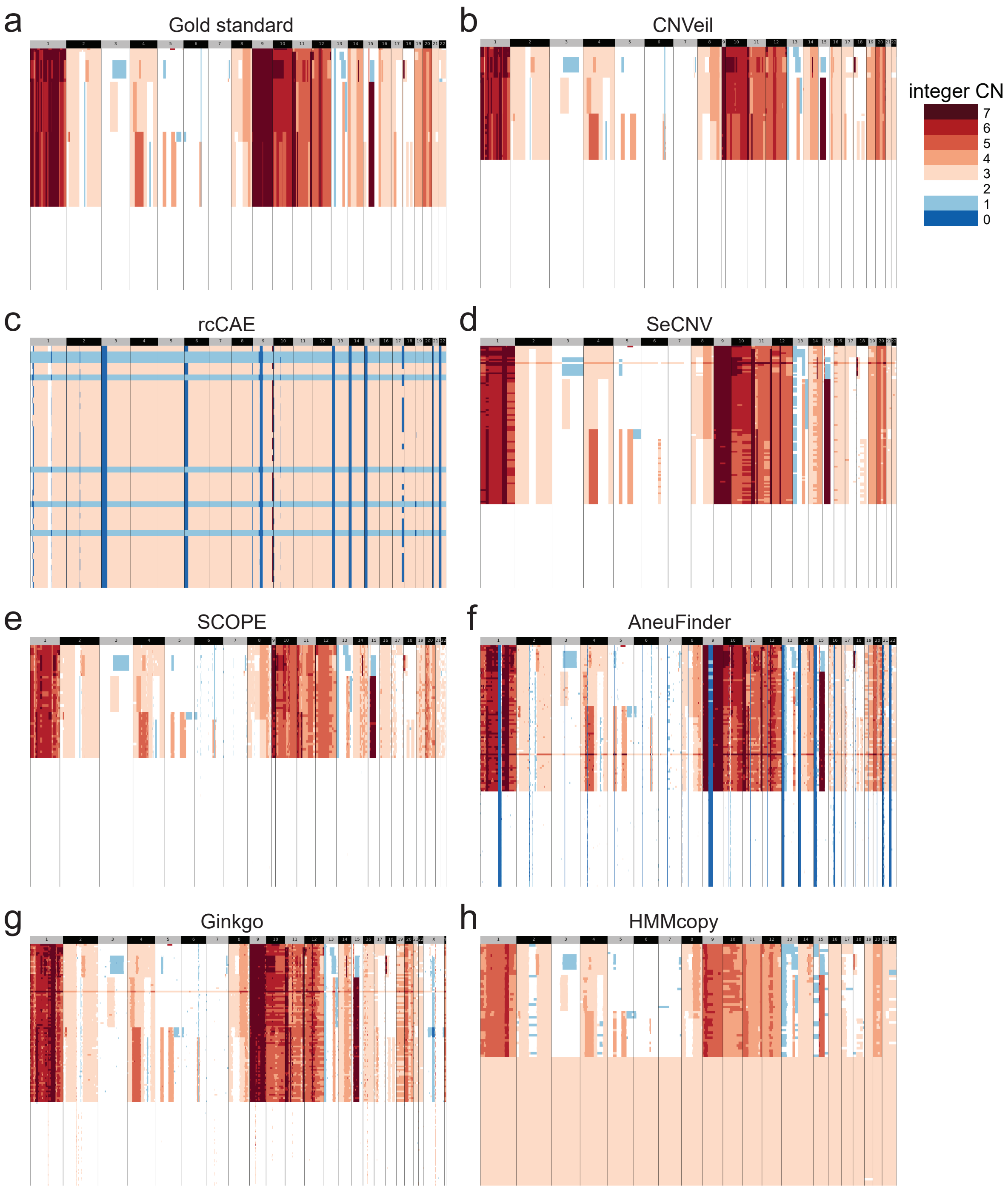

Figure S3: **Comparison of inferred copy number profiles of single cells from the simulated dataset with a ploidy of 4.0. (a)** Established copy number profiles by the gold standard. **(b-h)** Inferred copy number profiles by CNVeil, rcCAE, SeCNV, SCOPE, AneuFinder, Ginkgo, and HMMcopy. We ordered all heatmaps in a consistent cell order aligned with the gold standard which includes a normal cell subclone and a tumor subclone. Tools are shown in chronological order by publication year.

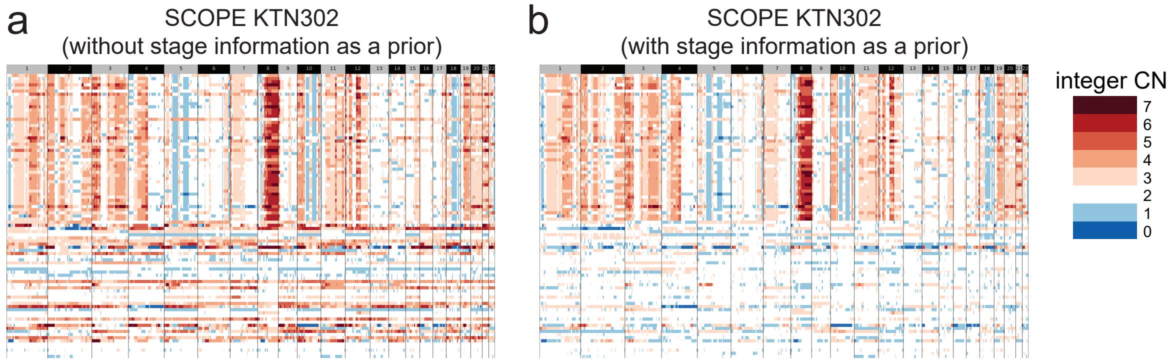

Figure S4: **Inferred copy number profiles of single cells from KTN302 by SCOPE with and without stage information as a prior.** **(a)** SCOPE employs the Gini coefficient to identify normal cells that act as the negative control; **(b)** SCOPE selects cells from mid-treatment stage cells as the negative control.
